# Mining Mass Spectra for Peptide Facts

**DOI:** 10.1101/2023.10.27.564468

**Authors:** Jeremie Zumer, Sebastien Lemieux

**Affiliations:** Institute for Research in Immunology and Cancer; DIRO, Université de Montréal, Montréal, Québec H3C 3J7, Canada; Department of Biochemistry, Université de Montréal, Montréal, Québec H3C 3J7, Canada

**Keywords:** Deep Learning, Spectrum Scoring, Peptide Identification, Proteomics, ProteomeTools

## Abstract

The current mainstream software for peptide-centric tandem mass spectrometry data analysis can be categorized as either database-driven, which rely on a library of mass spectra to identify the peptide associated with novel query spectra, or de novo sequencing-based, which aim to find the entire peptide sequence by relying only on the query mass spectrum. While the first paradigm currently produces state-of-the-art results in peptide identification tasks, it does not inherently make use of information present in the query mass spectrum itself to refine identifications. Meanwhile, de novo approaches attempt to solve a complex problem in one go, without any search space constraints in the general case, leading to comparatively poor results. In this paper, we decompose the de novo problem into putatively easier subproblems, and we show that peptide identification rates of database-driven methods may be improved in terms of peptide identification rate by solving one such subsproblem without requiring a solution for the complete de novo task. We demonstrate this using a de novo peptide length prediction task as the chosen subproblem. As a first prototype, we show that a deep learning-based length prediction model increases peptide identification rates in the ProteomeTools dataset as part of an Pepid-based identification pipeline. Using the predicted information to better rank the candidates, we show that combining ideas from the two paradigms produces clear benefits in this setting. We propose that the next generation of peptide-centric tandem mass spectrometry identification methods should combine elements of these paradigms by mining facts “de novo; about the peptide represented in a spectrum, while simultaneously limiting the search space with a peptide candidates database.

## Introduction

Mass spectrometry (MS) is currently the only high-throughput method that allows the analysis of peptides and proteins at scale^1^. Tandem MS is typically needed to accurately identify the fragments under study as mass conflicts make precursor fragment masses an insufficient proxy for peptide identity in peptide-centric experiments. In this paper, we focus on the tandem MS paradigm and use the term “mass spectrometry’ to refer to “data-dependent tandem mass-spectrometry’ (in particular in the peptide-centric identification setting) for the sake of brevity. Due to the large and complex data generated by mass spectrometry experiments, analysis by computer software of the mass spectra output is necessary to process the data at a reasonable rate. The main paradigms for such analysis are:

- de novo, or tag-based approaches, where the software tries to identify parts, or complete, sequences from the spectrum^2^;
- correlation-based approaches^3^, which attempt to describe the strength of the correspondence between an experimental spectrum and a spectrum from a database, possibly generated based on a candidate sequence^4^, and
- probabilistic models, which compute the probability that a certain amount of observed peak matches between an experimental spectrum and an expected spectrum (usually generated from a database) occurs by chance as a proxy for the likelihood of a match between the experimental spectrum and the peptide represented by the database spectrum^5^

Some of the most widely used mass spectrometry analysis software tools are proprietary. Vhile the high-level ideas behind these methods are sometimes published^3,5,6^, their finer-grained details remain trade secrets^5-7^, which makes iterating or probing these methods complicated. This means that for algorithm development and comparisons, only open-source software is viable (although this is not sufficient, as the design of the tool in question may make the integration of some algorithms hard or impossible). In many contexts, however, closed source tools are preferred for use in practice for peptide identifications^3,5^. Nevertheless, these proprietary methods see wide use as practitioners empirically find them to work best for their usual workloads, as reported in the literature^8,9^. Many open-source solutions are available (MS-GF+^10^, Comet^11^ or Morpheus^12^ for example), but they don t always respond to the community s expectations, often producing results of insufficient quality, although the situation may be changing. For example, IdentiPy^13^ was recently proposed as a new open-source algorithm that performs better than its other open-source peers, representing a further step toward closing the gap between open-source and proprietary methods.

Even with a wet lab protocol that is optimally tuned for a given (preselected) downstream analysis tool, the results obtained leave space for further improvement on the basis of correct peptide identification rate^1,14,15^. In addition, the various state of the art methods share a notable flaw: they rely heavily on input databases (and their specific contents, entry count, composition, and so forth) to query against candidates^l,5-7^, which makes results highly dependent on the input database design. In applications where a large specialized or extended database might be useful, the mainstream algorithms place technical restrictions on the sit’e of the databases vs correctly identified peptides, leading the community to develop various sub-optimal tricks to work around these issues^16^. Recently, to address the above, the Pepid project (https://github.com/lemieux-lab/pepid) has attempted to build a platform designed specifically for experimenting with peptide search algorithms, which may accelerate improvements in open-source search engines and to the development of more robust identification pipelines.

De novo sequencing methods are tandem mass spectrometry analysis methods that do not make use of databases, instead relying purely on the mass spectrum. Solving de novo sequencing has been attempted for decades with limited success^2,17-21^. However, some of these methods optimize metrics that implicitly imply good performance on other tasks: the recent method Deep Novo^2^, for example, computes amino acid-level errors over length of ground truth peptide, and errors over length of predicted peptide, in their experiments. Such a metric is consistent with the model s ability to predict peptide length. This suggests that the auxiliary peptide length prediction task may both be easier than, and a required component of, de novo sequencing. Moreover, the fact that DeepNovo performs well for the aforementioned metric, which implicitly measures length prediction performance (despite optimizing for de novo sequencing outcome), strongly suggests that peptide length can be predicted from just the spectrum.

There has been attempts to use de novo sequencing outputs to further improve database searches^3^, though those attempts have lead to unconvincing results in the general case, probably in no small part due to the disappointing performance of de novo sequencing methods when used in this context^2,17-21^. On the other hand, de novo sequencing methods have shown successful uses in constrained applications on clean spectra^22^. This demonstrates that de novo methods are sensible as a class of approaches, though hampered by the complexity of spectra from peptides of unconstrained structures. To the best of our knowledge, there has been no attempt so far to decompose the de novo problem into distinct subproblems (which could be easier than full, end-to-end de novo sequencing, and therefore achieve the subproblem s objective better than de novo approaches can perform end-to-end) and to use solutions to those subproblems for database-driven methods. In this work, we propose one such decomposition. Ve then focus on peptide length prediction as a subtask of the full de novo sequencing problem to demonstrate its potential usefulness in database-driven peptide identification. Previously, software like Max(2uant^23^ have modeled the relation between peptide length and scores generated by methods such as Mascot^6^, and to derive more confident identifications using that model. This is different from our methodology, where we propose to model the length of the peptide based on the spectrum only, and combine that with an external scoring method.

While there can be many ways to consider a decomposition for de novo sequencing, we choose the following model:

1. Peptide length prediction
2. Amino acid composition prediction
3. Amino acid ordering

We show that solving one subproblem from this example decomposition (namely, peptide length prediction) is both reliably achievable using deep learning, as well as proving a valuable asset as part of a state-of-the-art database-driven peptide identification pipeline.

In this paper, we demonstrate that a deep learning-based method can accurately predict peptide length from the spectrum, and that the predictions from such a method can further be used to achieve higher peptide identification rates at a fixed false discovery rate (FDR) as well as at fixed false discovery proportions (DP) using the SPOT^24^ peptide dataset ProteomeTools^15^.

## Methods

In order to establish a proper evaluation of the proposed methods, we make use of ProteomeTools^15^, a dataset of synthetic (SPOT^24^) peptides. This gives us access to a reasonable ground-truth for the peptide identities in the dataset as well as enabling FDP evaluation rather than relying on target-decoy FDR estimation. The availability of ground-truth data also allows a richer performance comparison between proposed and competing methods. The length model is trained on the Massive-KB^25^ dataset, so as to keep the evaluation data and the train data as separate as possible to show the applicability of our method. To further demonstrate the generalitzation power of our length prediction method, we also present measures on the One Hour Yeast Proteome^26^ dataset.

We present two measures of the effectiveness of our proposed approach:

1. Identifications under False Discovery Proportion (FDP)
2. Identifications under Target-decoy approach-based FDR (TDA-FDR)

Since we have access to reasonable ground-truth, we can compute FDP (sometimes called factual FDR^27^), which is not usually available in non-synthetic datasets where the identity of the peptides in a pool is not known. We compare the relative results (i.e. between the ground-truth and our proposed method) using the aforementioned metrics and note that due to the availability of ground-truth data, the identities at various FDP thresholds is most likely the most relevant. The FDR metrics serve as reference and offer a different, more common view of the results.

## Data

ProteomeTools is a dataset of spectra obtained with an Orbitrap Fusion Lumos on a pool of synthetic peptides determined by *in silico* trypsic digestion of the 42 164 sequences of human proteins found in the Swiss-Prot database at the time of recovery of the release in this paper. ProteomeTools data is available on the PRIDE archive with accession ID PXD004732^15^.

In the original ProteomeTools paper, spectrum quality was assessed using the Andromeda score^5^ for the ground-truth peptide (if it was identified at all by Andromeda). We show statistics related to spectrum quality in Figure 1. The distribution of masses vs peptide length with a linear fit in Figure 1A gives an idea of task difficulty. In Figure 1B, we show the distribution of peptide lengths across the dataset. Figure 1C and D show score distributions, where an Andromeda score of less than 100 is considered “low quality’ for the spectrum^5,15^. It can be seen that while there are many low-quality spectra, the majority are of relatively good quality. We used only the data from the higher-energy collision dissociation (HCD) experiments at a 25 normalized collision energy (NCE) as these seem to produce the best scored results out of the available data according to the analysis in the original ProteomeTools paper^15^. The length distribution of peptides in the dataset (fig. 1B) shows a preference for small peptides with 44.28% of the spectra in the dataset covering peptides shorter than 12 amino acids (which corresponds to 38.54% of unique peptide sequences).

**Figure 1:**
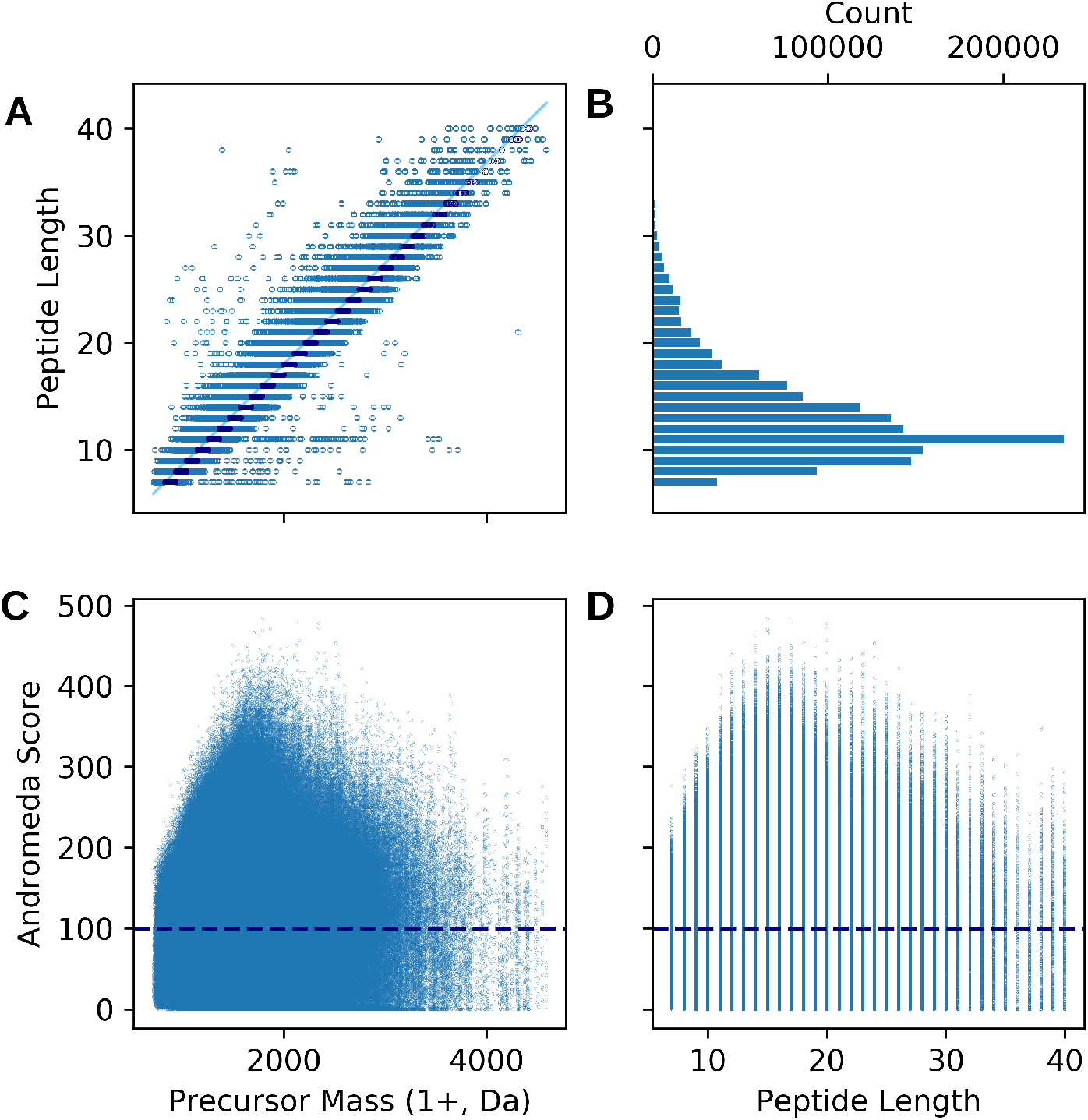
ProteomeTools dataset properties. **A:** Distribution of peptide lengths vs precursor masses in the dataset. The line represents a linear regression. Dark markers are precursor masses from the spectra with the correct length label using this linear regression, lighter markers are incorrectly labeled. **B:** Distribution of peptide lengths in the dataset showing a large proportion of shorter peptides. **C:** Precursor masses vs Andromeda scores. 52.44% of peptide-spectrum pairs pass the suggested quality threshold of 100 (as suggested by the Andromeda developers^5^), shown by the dashed line on the graph. **D:** Distribution of peptide lengths to Andromeda scores.

The One Hour Yeast Proteome is a dataset of non-synthetic yeast peptides with a baseline generated by the Thermo Fisher version of the SE(2UEST search engine^26^. Ve use the “batched’ dataset, which combines the result of all the hour-long runs. URLs to the Chorus Project submissions of the data are available in the One Hour Yeast Proteome paper^26^.

The Massive-KB dataset builds consensi based on various proteomics experiments submitted to the project, thus combining more realistic peptide pool searches with reasonably confident ground truth identities^25^. It is the largest of the 3 datasets (about 4x the size of the ProteomeTools dataset) and contains HCD-fragmented spectra of human peptides. The data may be obtained at https://massive.ucsd.edu/ProteoSAFe/static/massive-kb-libraries.jsp

To use Massive-KB for evaluation by the length prediction combination with Pepid, a subset of the data where the ground truth peptides may only be modified with fixed modification C(CAM) and variable modification M(ox) was used (by correspondence with ProteomeTools, and due to resource constraints).

In the same setting, the performance for the yeast dataset was obtained by performing the search only on the subset of the data with a ground truth identification (representing about 50% of the dataset) and uses SEQUEST search results rather than known real identities (hence the high accuracy vs pepid).

To ensure that the reported performance metrics generalize to never-before-seen peptides and spectra, the Massive-KB dataset, which is used for training, was split three-way according to the following criteria:

1. Data (spectra, spectrum metadata, associated peptide) is split into groups depending on the peptide they represent, with each group corresponding to one unique peptide sequence;
2. The collection of groups is then split 3-ways into sets: 80% for training, 10% for validation, and 10% for testing.

In total, the Massive-KB dataset we used contained 3 178 174 peptides, forming a training set of 2 542 539 peptides, and 317 817 peptides each for testing and validation. All models are trained using the training set, whereas the validation set is used to optimize the model choice and hyperparameters. Finally, the held-out test set is used to report all results in this work for length prediction performance. For experiments combining length prediction with pepid, the Massive-KB, ProteomeTools or One Hour Yeast Proteome dataset are used, as and where indicated, with the ProteomeTools and Yeast data serving as fully held-out data.

## Model

We developed a deep learning approach using combined multiple modalities to process the spectrum and metadata, as illustrated in Figure 2A. The mass is encoded as a simple number, while the spectrum is encoding as a rasterized vector of fixed sit’e 50000, summing in each vector element *v*_*i*_ the intensity of all peaks in the spectrum of m/z *i \** 10 to (*i* + 1) *\** 10. The model includes a fully connected network, a convolutional neural network, and a different network integrating metadata such as the mass in the captured spectrum (Figure 2B). At this stage, latent representations of those modalities are extracted by each network. Ve train an auxiliary length prediction task following each of these latent representations (Figure 2D). In Figure 2C, the different networks outputs are combined using another deep learning network. This network is trained independently of the subnetworks that provide modality-specific latent representations, and gradients do not flow below it: the subnetworks are trained using only the signal from their auxiliary training networks, with a different, independent predictive subnetwork head per modality, as depicted in Figure 2E.

**Figure 2:**
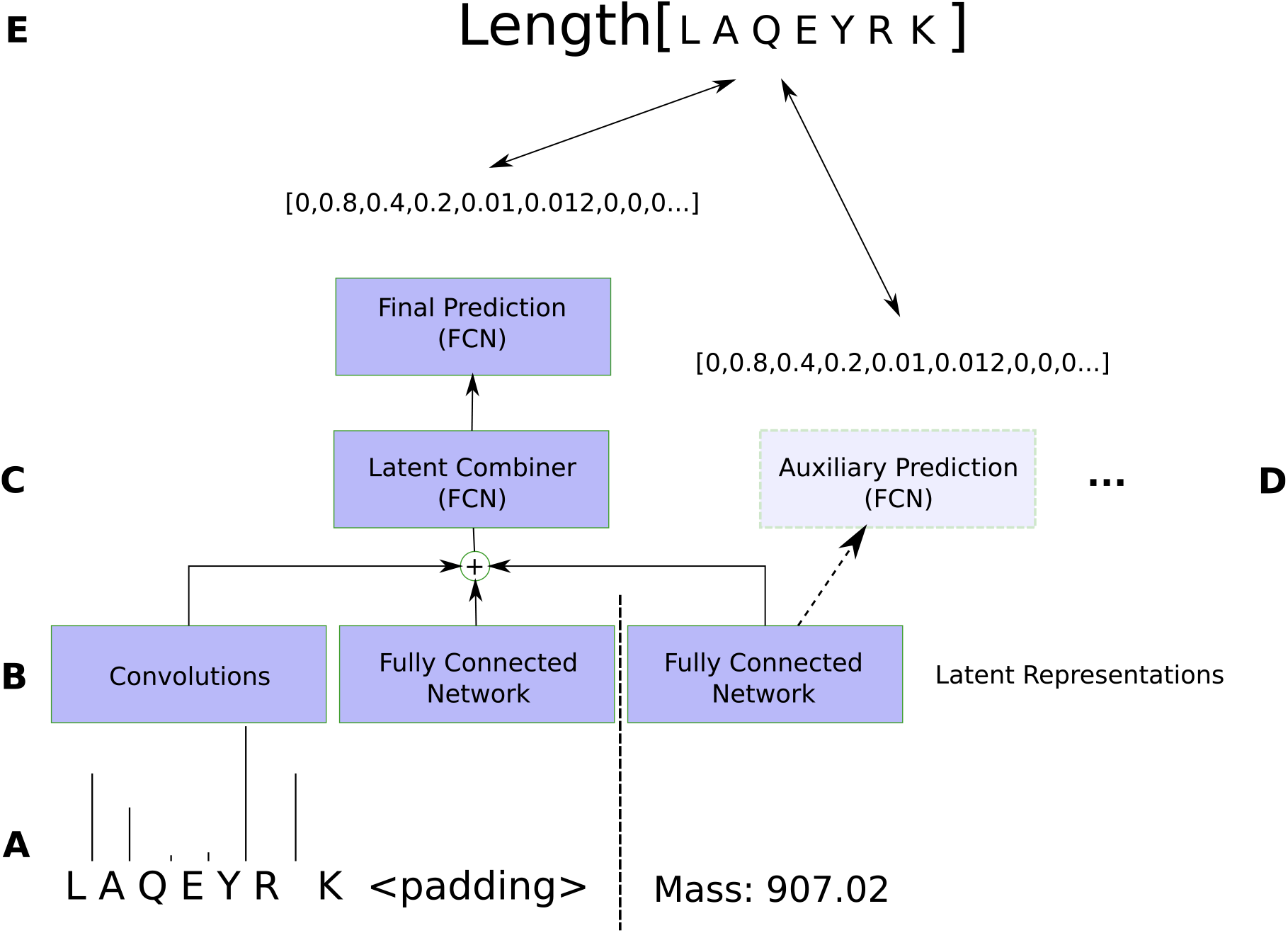
Diagram representation of the proposed length prediction model. **A:** Inputs. B: Modality-specific networks. **C:** Combiner network. **D:** Auxiliary prediction networks (training only). **E:** Prediction task and training feedback. See text for details.

The optimization algorithm is Adam^28^. When the validation loss does not achieve a new minimum in 10 epochs, the learning rate is divided by 10 and the best performing model is loaded to continue training with this new learning rate. This method is iterated 3 times starting at a learning rate of 10^*−*3^, after which no further improvement on the held-out validation set could be observed. The optimization problem is described as follows:

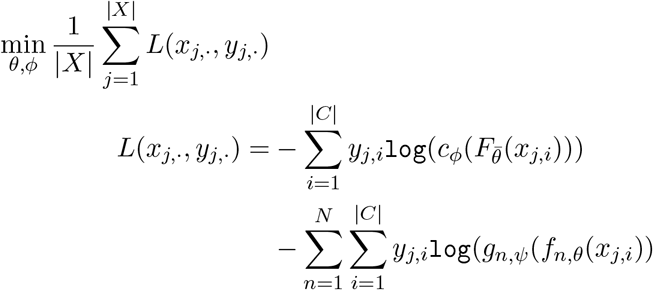

where *X ∋*(*x*_*i*_, *y*_*i*_ *∈ ℛ*^|*C*|^) is the training data, |*X* | is its size and |*C* | = 40 *−* 6 length categories are considered i.e. the model classifies between classes and, inclusively, using cross entropy loss. Optimization is performed over hyperparameters *ψ* for each prediction network used to help drive the subnetworks, *θ* which are the subnetwork parameters, and *ϕ* which are the combiner network’s hyperparameters. The neural networks *f*_*i*_ are the subnetwork ‘processors’, *F* is the concatenation of those networks, *N* is the count of subnetworks 3 in this case. *c* is the combiner network, and *g*_*i*_ is the subnetwork prediction output network for subnetwork *f*_*i*_. Additionally, we use the notation 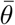 to represent *θ* with gradients disabled, i.e. interpreted as a constant. The full hyperparameters are described in Supplementary Material.

### Search Engine Integration

To show that length prediction can improve real peptide search result metrics, we integrate the score from peptide search using Pepid with our length prediction model by producing several features for rescoring by Percolator, as described later in the text. We chose Pepid because its flexible design, good wallclock performance, and high baseline identification performance allowed us to quickly implement the integration and to iterate on feature design for Percolator. This provides a realistic look at what length prediction can achieve in real-world search scenarios.

To showcase the advantage of length prediction in a familiar setting, we compute statistics of the length prediction namely the probability output from the model for the candidate’s length given the spectrum, the difference between this probability and the best-scoring, next-best scoring, previous-best scoring and worst scoring lengths, and the absolute and non-absolute difference between the best-scoring predicted length and the candidate length and provide them to Percolator ^29^ for rescoring.

## Results

We assessed the feasibility of the length prediction task by looking at the distribution of distinct lengths vs the mass of precursors in the dataset, which is shown in Figure 1A, clearly showing that the relation is linear but the range of possible lengths for any mass is wide. To establish a baseline accuracy, we used a simple linear regression, which was trained and tested on the entire dataset i.e. performance is reported on the same dataset used for training, achieving an accuracy of 3. 3. This suggests the task is not trivial and that methods based on precursor mass and peak-matching-based scores may not be able to exploit implicit peptide length data. We then used a linear regression baseline fit on the Massive-data to assess performance reusing the full set already used for fitting as an indication of performance upper bound, with results as per Table 1. We note that the performance on the Yeast and ProteomeTools datasets are higher than on Massive-, despite the model being trained on the same Massive-subset that it is being tested on. This is likely due to Massive-being the more realistic of the three, as the ground truth for the Yeast dataset is actually Sequest predictions, while ProteomeTools is a dataset of synthetic peptides.

**Table 1:** Linear regression baseline performance (trained on Massive-KB, tested on each dataset). MAE: Mean Absolute Error.

|  | Accuracy (%) | MAE |
| --- | --- | --- |
| ProteomeTools | 40.8 | 0.88 |
| Massive | 30.4 | 1.31 |
| Yeast | 42.0 | 0.83 |

To demonstrate that length prediction can positively impact peptide identifications beyond state-of-the-art methods, we show in Figure 3 that the Pepid search engine with artificial perfect length predictions generated from the ProteomeTools groundtruth followed by rescoring by Percolator ^29^ can consistently achieve higher identifications across common false discovery ranges. We also show that this holds for FDP i.e. the real metric of interest .

**Figure 3:**
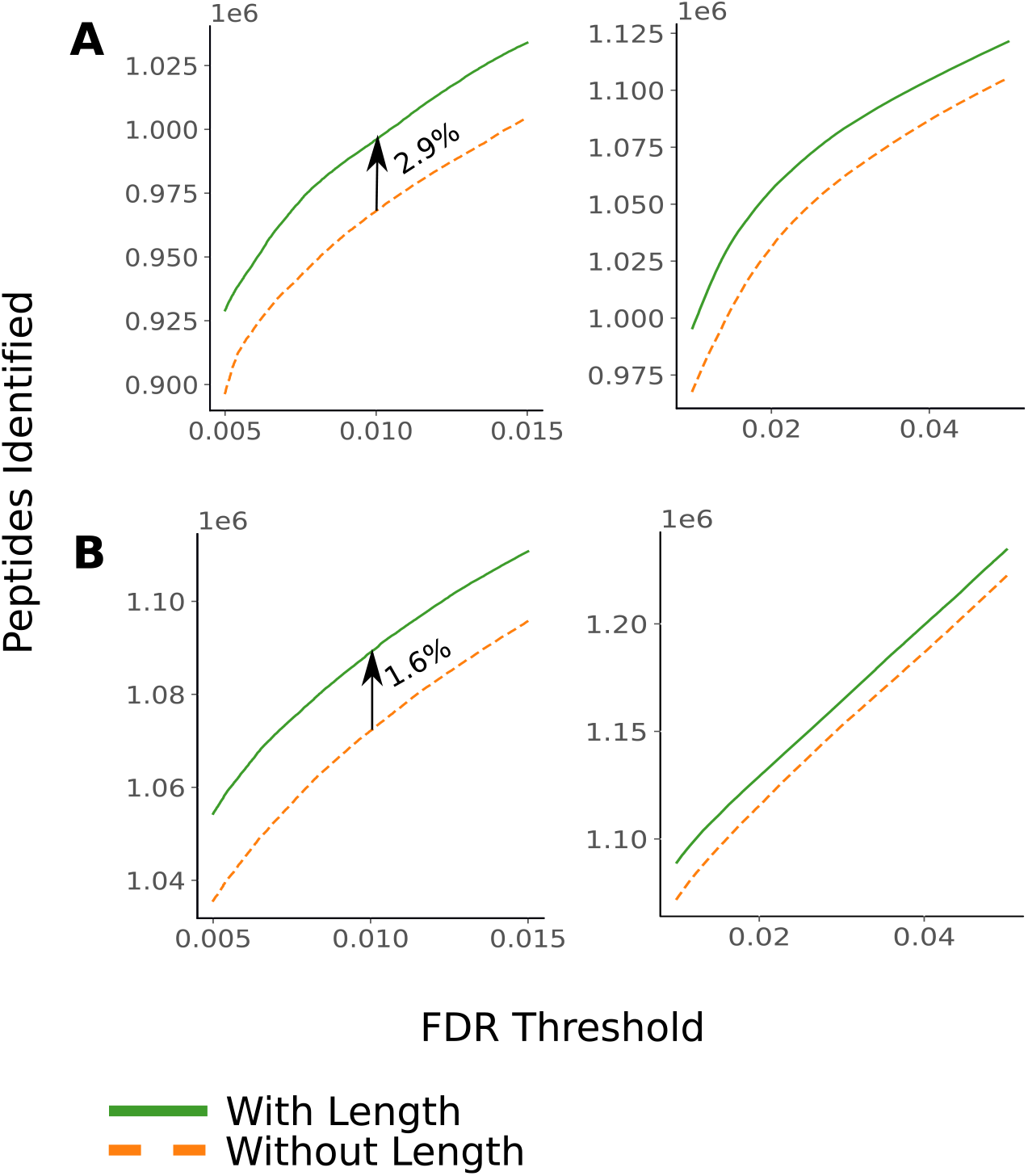
Pepid with a peptide length “oracle” (values obtained from the ground truth peptides in ProteomeTools). **A:** Identifications across false discovery proportions. **B:** Identifications under target-decoy-based false discovery thresholds.

The results show a modest, but consistent, improvement across the board using just Percolator. Combination using other strategies, such as candidate filtering, using more sophisticated, non-linear rescoring methods, ad hoc scoring functions that taken the length prediction into account, and so on, could potentially result in further improvements even with the oracle, although it is as yet unclear which of these methods might be most suitable, and how to best combine and prove them for next generation proteomics peptide analysis.

The length prediction model’s performance is presented as a confusion matrix in Figure 4. Since this task, to the best of our knowledge, has never been attempted before, we use the Pepid search engine’s results as a baseline by taking the length of the best-scoring PSM for each spectrum as a proxy for the “predicted length; from Pepid to be compared to our length prediction model’s output. In Table 2, prediction accuracy for the model trained on the Massive-model is compared based on the range of length of peptide sequences in amino acid, using either the dataset’s, or the pepid top-ranking predictions, as ground truth sequences from which the length is computed.

**Figure 4:**
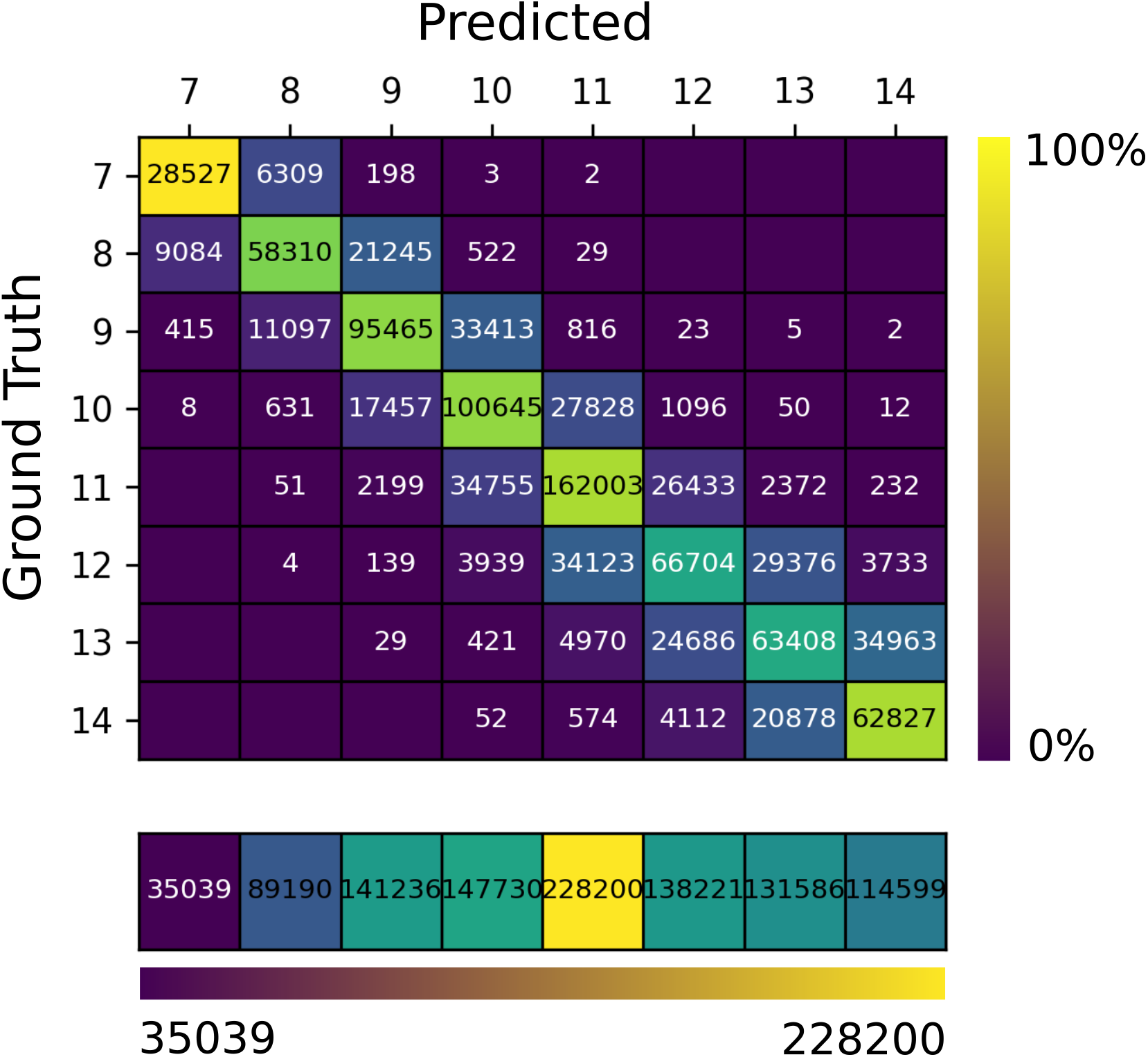
Length prediction model’s confusion matrix showing the total count predicted vs actual ground truth length from the ProteomeTools set. Only lengths 7-14 (corresponding to the most represented classes in Pro-ProteomeTools) are shown for easier viewing. Colors in the matrix are for percentage relative to ground truth counts (green is higher, blue is lower). The bottom row displays marginals on all the classes, with coloring as per absolute counts.

**Table 2:** Length prediction accuracy (percentage of correctly predicted peptide lengths).

| Length Range | 6-40 |  | 7-14 |  |
| --- | --- | --- | --- | --- |
| Ground Truth | Data | Pepid | Data | Pepid |
| ProteomeTools | 42.7 | 40.4 | 52.8 | 50.6 |
| Massive | 47.7 | 35.8 | 65.1 | 51.5 |
| Yeast | 44.2 | 43.2 | 54.2 | 53.6 |

We note that the deep learning model works significantly better than the baseline linear regression on Massive-compare Table l and Table 2, despite only using a disparate subset when testing the deep learning model, unlike in the linear regression case which presents results obtained on the same set used during fitting, despite showing more modest improvements for ProteomeTools and the Yeast dataset. As noted previously, we attribute these results to the lack of a real ground truth in the yeast dataset, and to the synthetic nature of the ProteomeTools data. Nevertheless, we find that the spectrum data clearly improves length prediction across the board. In particular, we note that the performance between the Pepid top-hit pepid uses an algorithm very similar to Sequest for all three predictions are quite closer between the linear regression and the length prediction model, than on the ground truth from the data. Since common search engines provide mass statistics to Percolator for rescoring, this may be a hint that Percolator can e effctively exploit mass - but not length - information to achieve improved identification rates.

To further see why the mass-based and spectrum-based length predictions differ, consider the consistent improvements obtained by providing length information to Percolator, both in oracular and in prediction forms. These results demonstrate that (1) real length information as in the oracle can help identifications quite a bit despite Percolator’s apparent ability to leverage mass to separate hits in rough length-like categories on account of the inherent correlation between mass and length), and (2) spectrum-based length prediction also provides information in excess of what Percolator can exploit using only its default features and masses.

Length prediction integration results are shown in Figure 5, demonstrating modest but consistent identification improvements across FDR thresholds similar, but less, than in the oracle version. These results show that the proposed deep learning model can already help improve peptide identification results, despite leaving space for further improvements. This is also reiterated in Table 3 which shows identification numbers under target-decoy (i.e. classic) and ground truth i.e. FDP based on dataset ground truth peptides) false discovery rates at select common threshold values.

**Figure 5:**
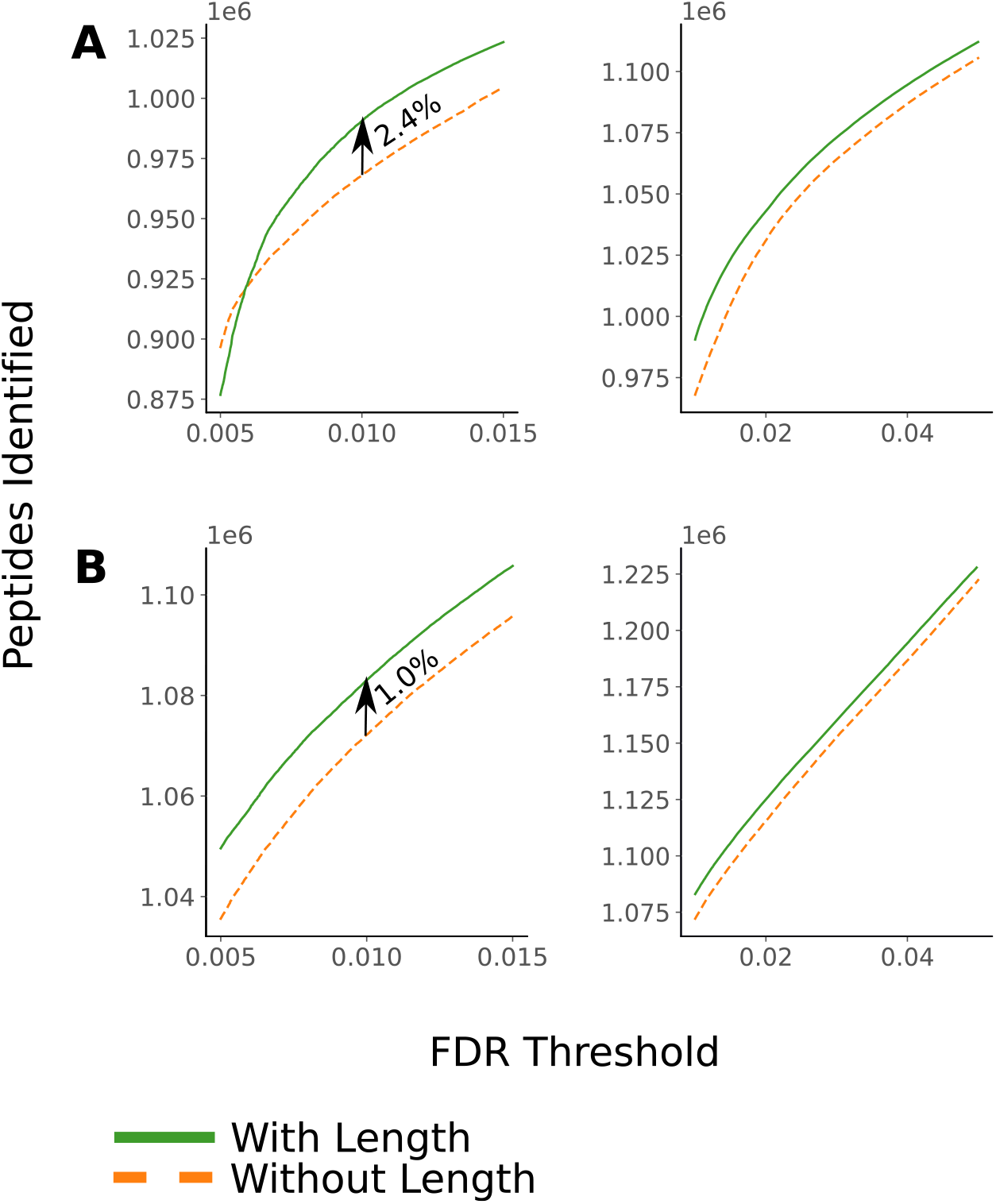
Pepid with the proposed length prediction model achieves improved performance compared to baseline. **A:** Identifications across false discovery proportions. **B:** Identifications under target-decoy-based false discovery thresholds.

**Table 3:** Performance results for the Pepid platform with and without the length model on ProteomeTools.

|  | ID @ 1% FDR | ID @ 1% FDP | ID @ 5% FDR | ID @ 5% FDP |
| --- | --- | --- | --- | --- |
| Pepid | 1 031 333 | 388 771 | 1 179 915 | 1 051 333 |
| + Percolator | 1 072 146 | 967 607 | 1 222 694 | 1 105 596 |
| + **Length** | 1 083 066 | 990 546 | 1 228 584 | 1 112 063 |
| Oracle | 1 089 048 | 995 584 | 1 235 215 | 1 121 710 |

We would also like to point out that while the improved identifications over target-decoy based FDR is advantageous, the more important metric is the increased identifications under false discovery proportion. As shown in our results, length prediction is significantly (around 2*×*) more impactful when considering true peptide identifications as opposed to target peptide identifications. Due to difficulties in obtaining robust ground truth peptide identities in practice, this metric is often overlooked in favor of the easier to compute TDA-FDR, yet we show here a divergence between the quality of the improvement under both metrics, showing that only considering TDA-FDR during the development and evaluation of refinements to peptide identification pipelines may be misleading. This observation may also impact the feature design for rescoring algorithm like Percolator, or the different score designs for common engines like Comet^11^ or X!Tandem^3^ better TDA-FDR identifications may hide lower FDP identifications ^27^.

## Conclusion

In this work, we have shown that mass spectra contain exploitable information beyond just serving as the basis for matching against a candidate database. We have proposed a deep learning-based method that can exploit query spectra to predict the length of the peptide which generated the fragmentation pattern, and we have demonstrated that this information can successfully be used to improve peptide identification at fixed FDP or FDR thresholds. We believe this is the first time it has been shown that peptide lengths can reliably be recovered directly from the spectrum. Our results were compared to the recent Pepid engine, a modern engine implementing the popular search algorithm Comet’s scoring function, demonstrating that length prediction can be of practical interest to improve peptide identification rates in real experiments.

In the typical FDR-based comparison setup, both using FDP and the more classical TDA-FDR estimation, our approach slightly but consistently increased the amount of identified peptides across a range of FDR values, and especially around the common lower FDR ranges (i.e. around 1%). When using FDP, our method similarly enables better separation of false vs true hits, causing an increase in the amount of correctly identified peptides over the baseline Pepid model.

The length-based model leaves room for improvement, achieving limited accuracy across the range of peptide lengths present in the ProteomeTools dataset. Despite this low performance on the task the model was trained for, its output can already be used to improve peptide identification, demonstrating that the proposed approach can unlock further improvements for peptide identification. Beside our oracle’s performance demonstrating what is possible at the upper bound with a much improved model, this also suggests that extracting more information from the spectrum rather than trying for full de novo sequencing, for example) may provide more tools to further improve peptide identification in database-driven peptide identification tasks perhaps properties like hydrophobicity, or the presence of specific amino acids in the sequence, can also be predicted and used in this manner. Furthermore, these approaches can create a toolbox of algorithms that could lead to similar improvements in a de novo peptide sequencing framework by greatly restricting the potential search space for peptide candidates.

Combining the length prediction from our model with a classical peaks-matching-based approach i.e. that of Pepid in our case) is a complex problem. We used a two-step approach in this work and not an end-to-end approach because we aimed to compare the contribution of our de novo subproblem model to an existing baseline following common modern identification workflow practices, and thus to minimize code and architecture changes to just those required to combine our method and the Pepid scoring method. In light of our encouraging results, we believe that an end-to-end scoring approach may further improve peptide identification compared to the results presented in this work. While we do not have the means to combine this method with some of the commonly used software like PeaksDB or MaxQuant’s Andromeda as they are proprietary, it would be interesting to verify that this method can improve performance when combined with them.

We believe that the above results clearly show that a mixed approach, i.e. combining ideas from the de novo sequencing and from the database-driven peptide identification, has a lot of potential to improve upon the current state-of-the-art. A database can be used to initially constrain the search space to a manageable subset, and information inherently present in the query spectrum can further be used to reduce this subset by more confidently ranking candidates based on additional information not present in peak matching alone. This proposed approach does not suffer from what is arguably de novo sequencing approaches’ biggest weakness: the combinatorial search space and increased error probability scaling with peptide length, while making use of their strength: the ability to mine information from the spectrum in advance. Beside peptide length from just the spectrum, data like peptide amino acid composition, or properties like hydrophobicity, could be extracted to further improve identification rates. In addition, predictions from the sequence candidates, such as spectrum prediction or retention time prediction, could further be used in tandem with these spectrum-based feature extractors to improve identifications even further.

Our code is available as open source software as an integrated package in the Pepid framework starting with version v1.1.0. The repository may be accessed at https://ithub.cm/emieux-ab/pepid.

## Acknowledgenent

The authors would like to thank Assya Trofimov and Mathieu Courcelles for fruitful discussions and support. This work was funded by a grant from the Institut de Valorisation des Données IVADO .

## Supplementary Material

### Length Confusion Matrix

**Supplementary Figure 1 1:**
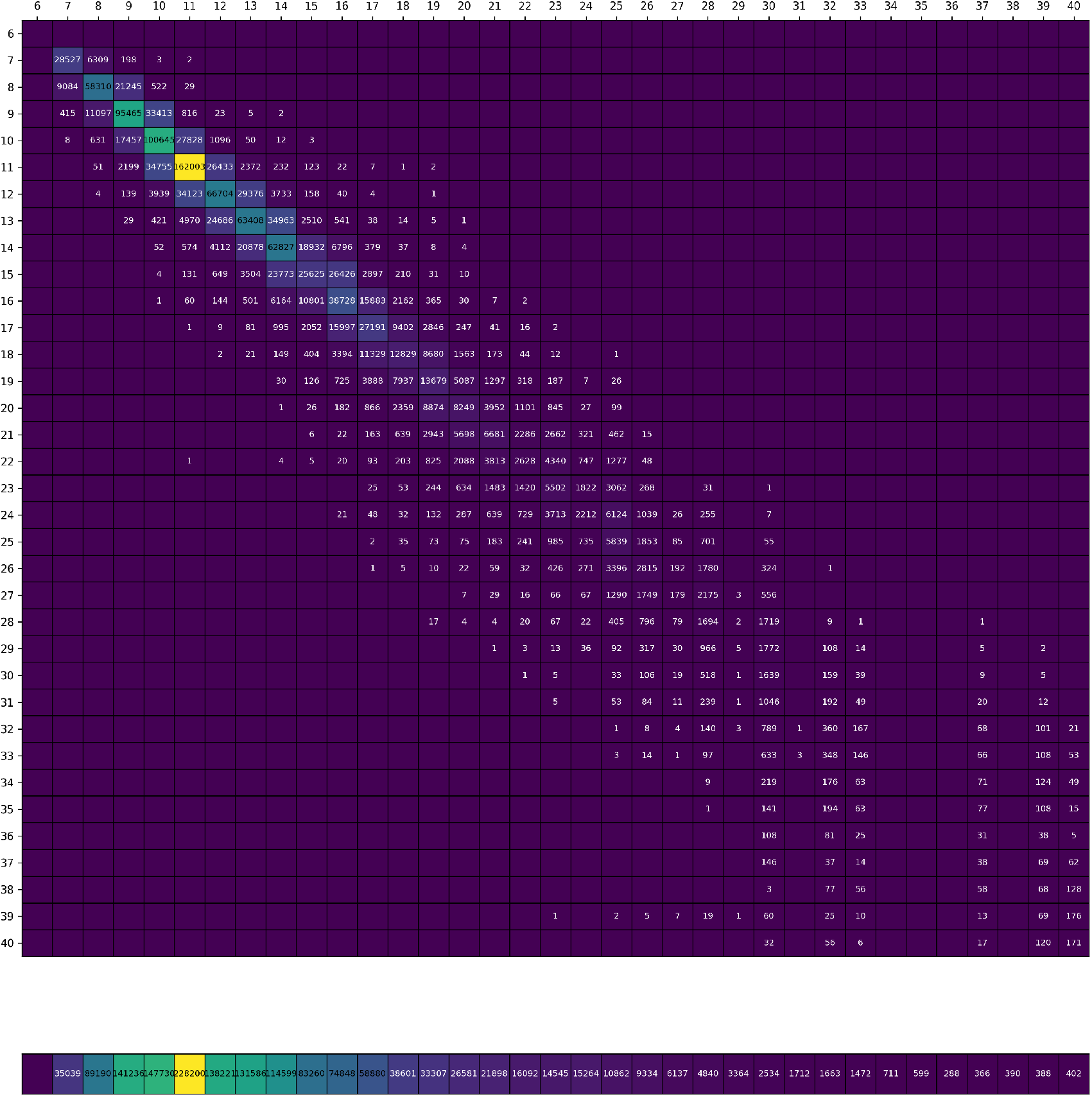
Full length prediction confusion matrix across the lengths the model was trained for (6-40 inclusive). The bottom row shows marginal counts for each class.

### Model Details

The model architecture is presented in the main text. Here we describe the exact hyperparameters in the final model.

The convolutional layers in the preprocessing stack were procedurally determined using the starting dimensions of the input. In total, 13 layers, composed of a convolution, a batch normalization (batchnorm), and rectified linear (ReLU), and a pooling layer. All pooling layers were mean-pools of size and stride 2. For the convolutional layers, they had a dimension of 11, unless the input size was not compatible, in which case they were 12. The last layer had a kernel size of 1. All layers used an embedding size of except the final layer, which converts the latent units into our output size of 3. The pattern of kernel sies was thus 11, 12, 11, 12, 12, 11, 12, 11, 12, 12, 11, 11, 1. The final layer is followed by a non-linearity of log-softmax rather than a ReLU.

To convert the convolution outputs to a latent representation compatible with the prediction “head;, a convolution of kernel size 1 and a pooling layer of size and stride 2 was applied.

The linear layers processing the spectrum are composed of fully-connected layers with units in their latent spaces. The input of the first network is the whole spectrum, of size A 9th fully-connected layer converts the latent output to our output size of 35 .

The linear metadata-processing network is composed of fully-connected layers with l units each, followed by one last layer with 3 outputs. The combination network has the same topology, but while the metadata network takes just the input mass as a feature, the combination network takes 11 features, namely the latent representations of the other networks as a concatenated vector.

## Notes

### Competing Interest Statement

The authors have declared no competing interest.

